# Structural co-modulation: An individualized measure of inter-component interactions in source-based morphometry

**DOI:** 10.64898/2026.01.26.701772

**Authors:** Aline Kotoski, Najme Soleimani, Sir-Lord Wiafe, Spencer Kinsey, Vince Calhoun

## Abstract

Source-based morphometry (SBM) is a powerful multivariate method for identifying covarying structural brain networks. However, standard SBM provides only a single loading value per component for each subject, which limits the characterization of relationships between these components. We propose a novel technical co-modulation approach to derive an individualized, network-like measure of structural brain organization. This method transforms the subject-specific SBM loading vector into a symmetric co-modulation matrix by computing the vector’s outer product. Each element of this matrix quantifies the pairwise interaction between structural components, creating a subject-specific fingerprint. Similar to functional connectivity that maps the temporal synchronization between networks, this matrix maps their joint structural prominence, reflecting how strongly two networks co-occur within an individual. To demonstrate the utility of this method, we applied it to structural MRI data from 210 patients with schizophrenia (SZ) and 195 healthy controls (HC) from the fBIRN psychosis dataset using functional networks as priors for SBM. We observed widespread reductions in structural co-modulation in the SZ group, particularly within and between visual, default-mode, and cognitive control networks. Furthermore, co-modulation patterns were significantly correlated with cognitive performance and clinical symptom severity in patients. Structural co-modulation provides a robust framework for quantifying individualized relationships between structural brain features, overcoming key limitations of standard SBM and offering a new avenue for integrating structural and functional brain analyses.

## 1. Introduction

Source-based morphometry (SBM) is a type of structural brain analysis focused on estimating maximally independent covarying patterns of, e.g., gray matter, that are individually expressed to different degrees (Xu et al., 2009). While SBM has successfully identified interpretable and biologically meaningful networks (Van Assche et al., 2024, Gupta et al., 2019, Jo et al., 2025), it provides subject-specific loading parameters (one per component) that can be subjected to statistical testing. This inherently limits the method’s ability to capture shared or interacting structural variations among networks that co-vary in a coordinated manner across individuals. To address this limitation, we propose a novel co-modulation approach that extends SBM by quantifying the degree to which different structural components co-vary across subjects, allowing for the investigation of higher-order statistical relationships between structural patterns. This framework provides a richer representation of structural brain organization, complementing conventional SBM by revealing inter-component relationships that may underlie large-scale coordination of structural changes. Here, we demonstrate that structural co-modulation is a highly sensitive marker for schizophrenia. To our knowledge, this is the first study to propose and implement structural co-modulation analysis of structural MRI (sMRI) features, providing a new avenue for exploring the interdependence of structural networks in both healthy and clinical populations.

Schizophrenia provides an important test case for evaluating the utility of the co-modulation framework, as it is a disorder characterized by widespread and heterogeneous brain abnormalities (van Erp et al., 2018). Numerous studies using voxel-based and source-based morphometry have identified gray matter reductions in distributed cortical and subcortical regions, including the frontal, temporal, and limbic areas, along with disruptions in large-scale functional and structural networks (van Erp et al., 2018, Zhou et al., 2025, Zalesky et al., 2012). These findings suggest that schizophrenia is associated not only with localized deficits but also with abnormal integration among brain systems. However, traditional structural analyses often focus on component-specific loading parameters, overlooking potential coordinated variations among regions that may reflect system-level pathology. Despite strong evidence for network-level dysfunction in schizophrenia, little is known about how structural networks interact or co-vary in this disorder. This has been studied at the study or group level by computing the structural network connectivity (SNC). However, this does not provide individual subject estimates (Alexander-Bloch et al., 2013). By capturing inter-component (i.e., inter-network) dependencies using an individual subject calculation, the co-modulation approach offers a new way to study whether the altered brain organization in schizophrenia extends beyond individual networks to encompass disrupted relationships among them.

Re-emphasizing the limitations of traditional methods like SBM in treating structural systems as isolated sources, other approaches have attempted to model structural relationships, most notably structural network covariance (SCN). However, SCN is typically computed at the group and cannot capture subject-specific alterations that may be clinically meaningful. This is a critical gap, as schizophrenia is increasingly recognized as a disorder of disrupted brain coordination rather than isolated regional deficits (Zhou et al., 2025, Gupta et al., 2019). The co-modulation framework directly addresses this limitation by modeling inter-component dependencies at the individual level, enabling the quantification of coordinated or dis-coordinated structural organization.

In this study, we introduce the co-modulation framework and apply it to an sMRI dataset of schizophrenia patients and healthy controls to demonstrate its utility. We first used this method assess group differences. We then investigated the clinical and cognitive significance of these patterns by correlating subject-specific co-modulation values with cognitive performance and clinical symptom scores. This technical note provides the first application of co-modulation, offering a novel and clinically relevant method for quantifying the interactive organization of structural brain networks.

## 2. Methods

Uniquely, the co-modulation framework extends traditional source-based morphometry by shifting the level of analysis from isolated components to their pairwise interactions. This transformation generates subjects-specific structural networks, allowing us to quantify coordinated organizational pattern rather than simple volumetric variations (Kotoski et al., 2024b). By transforming each subject’s SBM loading vector into a co-modulation matrix through outer-product computation, we can evaluate pairwise relationships between components at a single subject level. This approach enables assessment of large-scale structural coupling patterns and facilitates integration with other modalities, such as functional network connectivity in fMRI, for multimodal analyses.

The co-modulation approach is built upon parameters derived from source-based morphometry, a multivariate data-driven method that applies ICA to gray matter maps to identify networks of structurally covarying brain regions across a group of participants (Xu et al., 2009). Each resulting SBM component represents a distinct spatial pattern of gray matter, and each subject is assigned a loading parameter that quantifies their individual expression of that component. Prior work has demonstrated that the spatial patterns identified by SBM often show a remarkable correspondence to well-known resting-state functional networks, such as the default mode network (Luo et al., 2012). This inherent link between structural covariance and functional organization motivates the use of functional network maps as spatial priors to guide the SBM decomposition, a strategy we leverage in the current study to ensure a direct correspondence between structural and functional domains.

The SBM loading parameters that serve as the input for co-modulation can be generated using several distinct strategies, each offering different advantages. A primary mode of operation is a fully data-driven SBM, where spatial components are estimated directly from the structural data itself, allowing the discovery of novel covariance patterns (Xu et al., 2009). A second approach utilizes predefined anatomical or functional atlases, where parameters are derived by averaging gray matter within specific regions of interest (Tzourio-Mazoyer et al., 2002). A third strategy, and the one we employ in this study, involves using group-level functional network maps, such as those derived from fMRI-based ICA as spatial priors for a spatially constrained SBM analysis (Luo et al., 2012).

We selected this third, functionally informed approach for the present technical note. By using well-established fMRI network maps as priors, we directly link the resulting structural co-modulation patterns to known functional brain systems, facilitating a more straightforward interpretation and cross-modal comparison. Figure 1 shows the workflow of this study.

**Figure 1.**
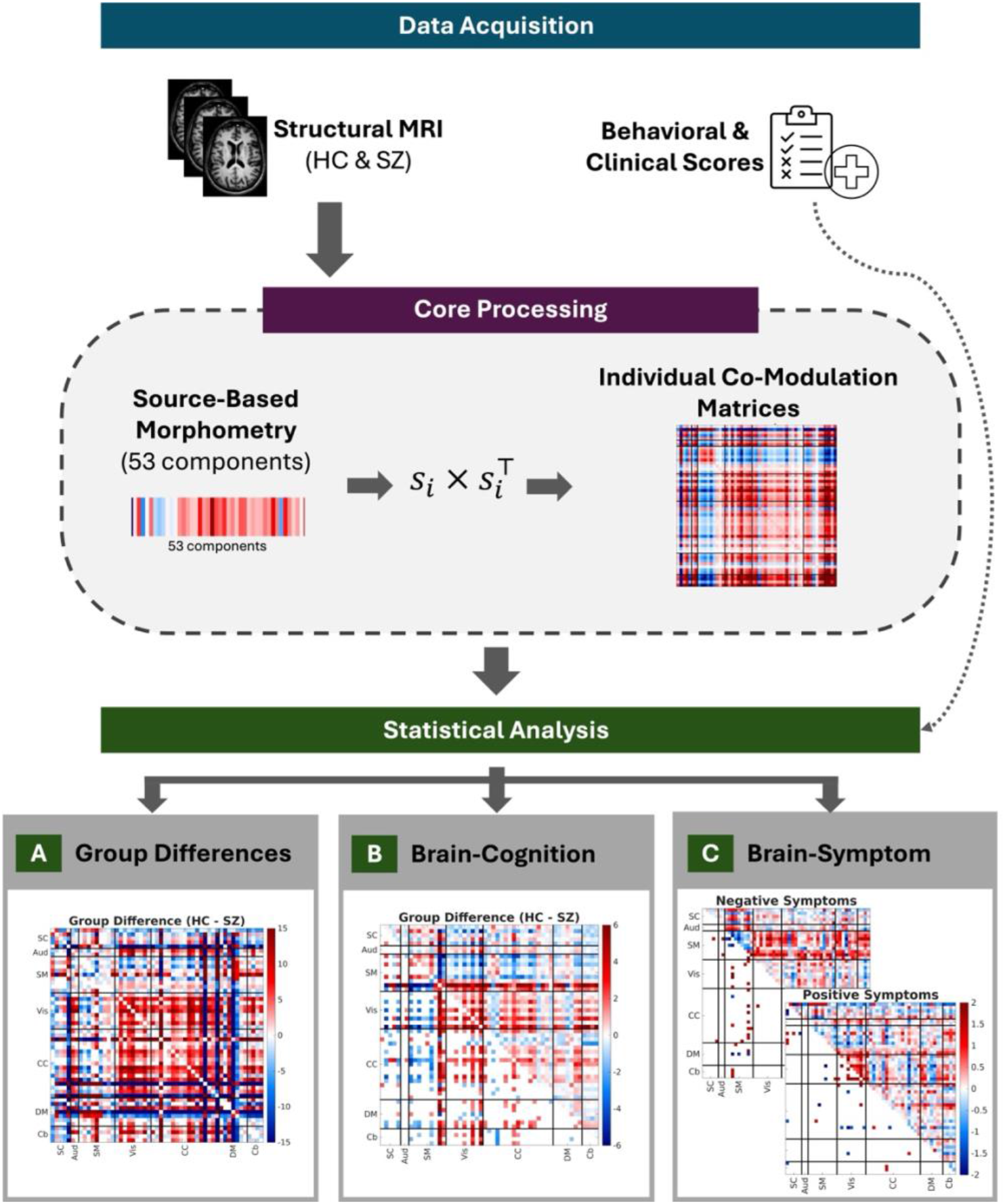
This diagram illustrates the sequential steps of the study, from data acquisition to the three primary modes of co-modulation application. Structural MRI data from healthy controls and schizophrenia patients were first processed using SBM. Subject-specific co-modulation matrices were then calculated. These matrices were subsequently used to investigate A) group differences in structural co-modulation between healthy controls and schizophrenia patients, B) relationships between co-modulation patterns and composite cognitive scores, and C) associations between co-modulation patterns and clinical symptom severity (positive and negative symptoms) in the schizophrenia group.

### 2.1. Participants

For this study, we used the Functional Imaging Biomedical Informatics Research Network (fBIRN) dataset., which comprises data collected as part of a multi-site brain imaging psychosis consortium. The schizophrenia group (SZ) consisted of 210 participants, aged between 18 and 62 years old (mean = 39.4 years; SD = 11.7 years). The healthy control group (HC) consisted of 195 participants, aged between 19 and 60 years old (mean = 37.7; SD = 11.2). With 161 males in the schizophrenia group and 139 males in the control group.

### 2.2. Imaging parameters and Preprocessing

Data were collected with two types of scanners (3T Siemens and 3T GE). The MRI sequences used for this work were a T1 structural scan (TR = 2.3/5.95 s, respectively; TE = 2.94/1.99 ms, respectively). The full details of the imaging acquisition protocol are described in (Turner et al., 2013).

Structural MRI data were preprocessed using the Statistical Parametric Mapping (SPM) software. Spatial normalization was performed using the DARTEL (Diffeomorphic Anatomical Registration Through Exponentiated Lie Algebra) algorithm to create a study-specific template and register images to MNI space with high precision. The resulting gray matter maps were modulated to preserve total tissue volume and resampled to a voxel size of 1.5 × 1.5 × 1.5 mm. Finally, the maps were smoothed with a 4 mm full width at half-maximum (FWHM) Gaussian kernel to reduce noise while preserving local anatomical detail.

### 2.3. Cognitive scores and symptoms

Clinical symptom severity in patients with schizophrenia was quantified using the Positive and Negative Syndrome Scale (PANSS) (Kay et al., 1987). The PANSS provides a comprehensive assessment of psychopathology. The positive subscale quantifies symptoms characterized by an excess or distortion of normal functions, such as hallucinations, while the negative subscale quantifies symptoms characterized by a reduction in normal function, such as blunted affect (Andreasen et al., 1999). For this analysis, we focused on the positive and negative symptom subscale scores. Cognitive performance was assessed in all participants using the Computerized Multiphasic Interactive Neurocognitive System (CMINDS ®) (O’Halloran et al., 2008). CMINDS is a battery of neuropsychological tests designed to measure performance across key cognitive domains, including processing speed, attention, working memory, and verbal and visual learning (O’Halloran et al., 2008). For this study, a composite score representing overall cognitive function was derived from the CMINDS battery for the correlation analyses.

### 2.4. Co-modulation

To implement the co-modulation approach, we first performed an SBM analysis using sMRI data, resulting in subject- and component-specific SBM loadings. Subject-specific SBM loadings represent the degree to which each individual expresses each structural component. In this study, we first calculated a spatially constrained ICA using the 53 components Neuromark_fMRI_1.0 template (Du et al., 2020) (Figure 2). The validity of using these functional priors to constrain structural decomposition has been demonstrated in our recent work, which showed that this approach captures robust structural covariance networks that align with established functional systems (Kotoski et al., 2024a). This results into a 53-dimensional vector of SBM loadings per subject.

**Figure 2.**
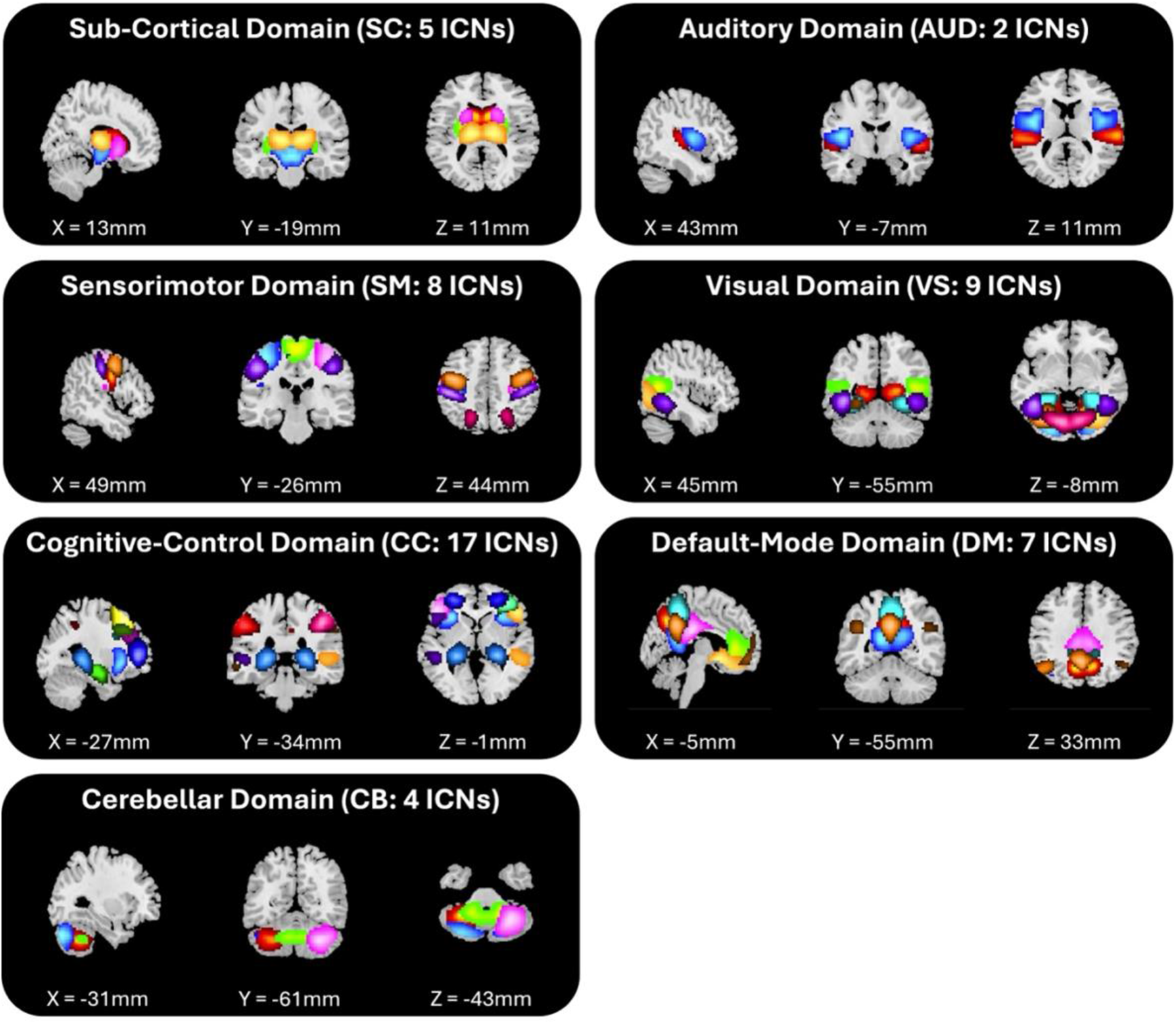
NeuroMark templates used to estimate 53 structural components. Intrinsic connectivity network (ICN) templates are visualized according to domain using an empirical threshold (Z = 1.96) with X, Y, and Z coordinated listed relative to the origin. Sevel displayed domains include sub-cortical (SC), auditory (AUD), sensorimotor (SM), visual (VS), cognitive-control (CC), default-mode (DM), and cerebellar (CB). Individual ICNs are displayed using distinct colors.

To quantify the relationship between structural components, we then computed the co-modulation matrix for each subject. For subject *i*, the 53-dimensional SBM loading vector (*s*_*i*_ ∈ ℝ^53×1^) was used to generate a symmetric co-modulation matrix (*M*_*i*_ ∈ ℝ^53×53^) defined as:

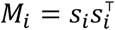

Each element *M*_*i*_(*j, k*) reflects the degree to which components *j* and *k* co-vary in their expression within that subject, capturing network-like relationships between structural sources. This computation produced one co-modulation matrix per participant for subsequent group-level analyses.

Conceptually, co-modulation extends beyond traditional SBM loadings by characterizing pairwise statistical interactions among structural component loadings, rather than treating them independently. While SBM identifies spatially distinct gray matter networks, co-modulation quantifies the extent to which these networks are jointly expressed, resulting into a subject-level matrix that is mathematically analogous to instantaneous temporal correlation of fMRI ICNs, which provides an estimate of the correlation at a single timepoint (Wiafe et al., 2026). Importantly, this formulation structural relationships between brain features enables direct comparison and integration of structural and functional data.

### 2.5. Application to schizophrenia

We performed three statistical analyses. Prior to all tests, the effects of age, sex, and scanning site were regressed out of the co-modulation values. First, element-wise group differences between HC and SZ co-modulation matrices were assessed using two-sample *t*-tests across subjects. A significance threshold of *p* < 0.05 was used, with corrections for multiple comparisons applied using the false discovery rate (FDR) method Benjamini-Hochberg (Benjamini and Hochberg, 1995).

Next, we investigated the relationship between brain co-modulation patterns and clinical symptom severity in patients with schizophrenia. Associations with both positive and negative symptom scores were analyzed. The Pearson correlation coefficient was computed between the symptom scores (positive and negative, separately) and the co-modulation value of each matrix element across all patients. A significance threshold of |*p*| < 0.05 was used, with multiple comparisons corrected using the false discovery rate (FDR) technique method Benjamini-Hochberg (Benjamini and Hochberg, 1995).

Finally, to study the brain co-modulation correlates with cognitive performance, we computed the Pearson correlation between cognitive scores and the co-modulation values for each matrix element and tested for significant relationships within each group. Additionally, we assessed group differences in brain-cognition relationships by computing a group difference correlation matrix and testing for significance. For all analyses (HC, SZ, and HC-SZ), a significance threshold of *p* < 0.05 was used, with multiple comparisons corrected using FDR method Benjamini-Hochberg (Benjamini and Hochberg, 1995).

## 3. Results

### 3.1. Co-modulation differences between schizophrenia and controls

For the co-modulation group analysis, we observed that the mean co-modulation matrix in the SZ group showed overall lower values compared to the HC group (Figure 3a). Visual inspection revealed higher co-modulation values in the HC group particularly within the visual and cognitive control domains, as well as between the cognitive control and visual domains, the default-mode and visual domains, and the default-mode and cognitive control domains. The mean absolute co-modulation value was 4.39×10−4 for the HC group and 3.76×10−4 for the SZ group (Cohen’s *d* = 0.83). To further isolate group-specific patterns, we visualized the group means after removing the average co-modulation across all participants (Figure 3c). These demeaned matrices highlight each group’s deviation from the grand mean. The HC group’s matrix shows prominent values above the grand mean, indicating stronger co-modulation than average particularly within and between the visual and cognitive control networks. Conversely, the SZ group’s matrix displays values below the grand mean, reflecting weaker co-modulation than average in these same domains.

**Figure 3.**
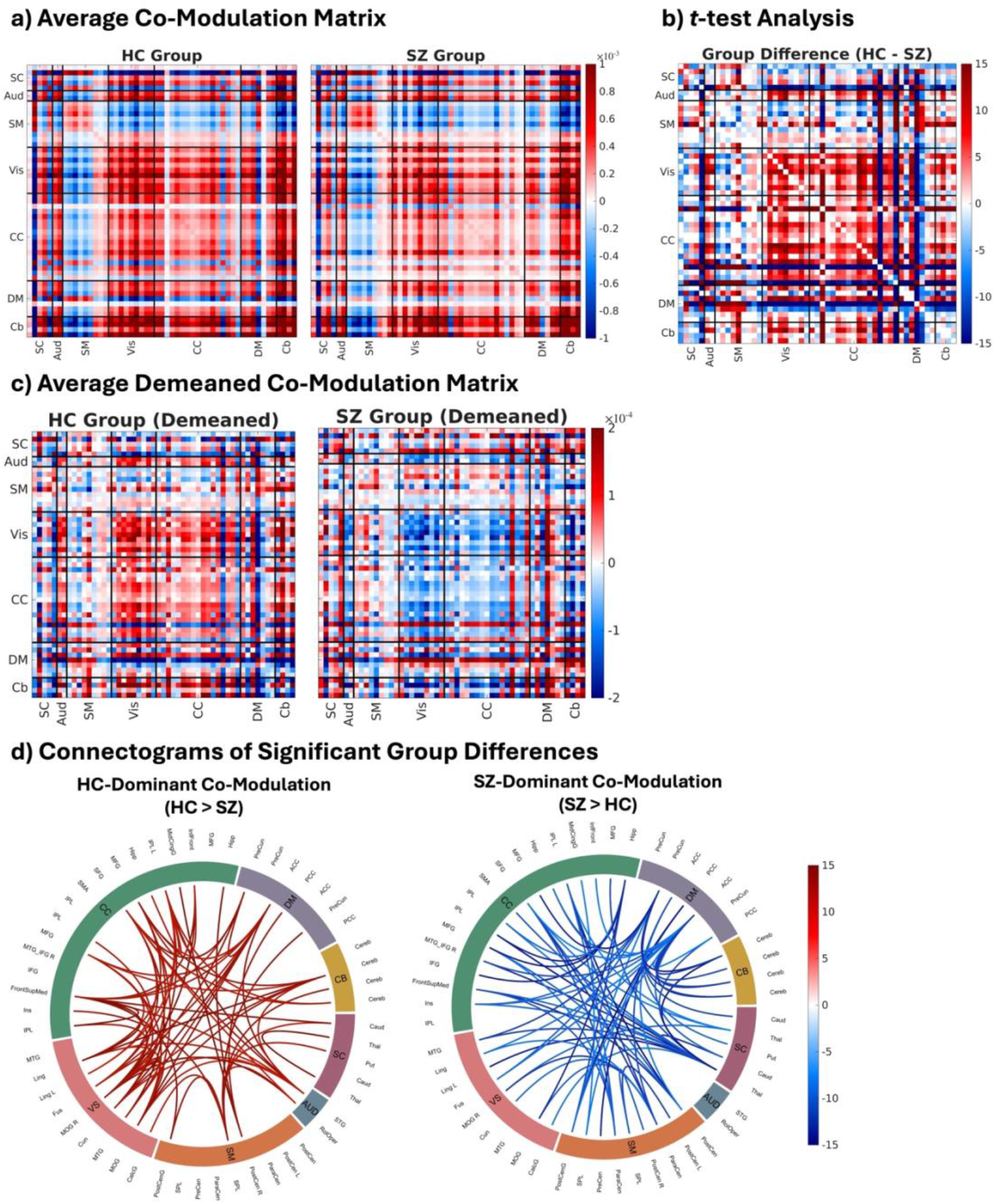
Group co-modulation matrices and differences between healthy control and schizophrenia. a. Mean co-modulation matrices for HC and SZ groups. The HC group exhibited higher co-modulation, particularly within visual and cognitive control domains, and between default-mode, visual, and cognitive control networks. b. Group differences (HC-SZ) assessed with a t-test. The group differences are shown as signed log_10_q (FDR corrected) values, positive values indicate HC > SZ, negative values indicate SZ > HC. Upper triangles display all edges, and lower triangles show only significant q-values, revealing widespread reductions in coordinated structural modulation in SZ. c. Demeaned co-modulation matrices, illustrating each group’s deviation from the grand mean. HC participants show predominantly positive deviations (values above the grand average), whereas SZ participants show predominantly negative deviations (values below the grand average), highlighting a global reduction in structural coupling in schizophrenia. d. Connectogram illustrating the top 5% most significant group differences. To visualize the specific nature of the disruption, results are separated by the direction of the contrast. The left panel displays connections where co-modulation was significantly stronger in controls (HC > SZ), and the right panel displays connections where co-modulation was significantly stronger in patients (SZ > HC). Line color represents the magnitude of the difference (-log10(q), FDR corrected).

Statistical tests for differences between cohorts revealed widespread differences in co-modulation across the brain (Figure 3b). The matrix displays the signed log_10_(q) values, where warm colors indicate higher co-modulation in controls (HC > SZ) and cool colors indicate higher co-modulation in schizophrenia (SZ > HC). The upper triangle shows all values across the full matrix, while the lower triangle displays only statistically significant edges after FDR correction (q < 0.05). Overall, controls exhibited stronger co-modulation across most domains, particularly within and between the visual, cognitive control, and default-mode networks, indicating a global reduction of coordinated structural modulation in schizophrenia. Biologically, this global reduction suggests a ‘structural decoupling’ of the brain’s architecture, implying that large-scale networks in schizophrenia fail to maintain the coordinated anatomical integrity and integration observed in healthy controls.

Connectograms reveal the extensive network of significant associations (Figure 3d). Given the high density of significant results, we are showing the top 5% most significant component associations and separated them into distinct plots based on the direction of the group difference to enhance clarity and interpretability. In both plots, the connections display the signed log_10_(q) values. The left panel displays connections where co-modulation was significantly stronger in controls, while the right panel displays connections where co-modulation was stronger in patients. Our analysis revealed a complex and widespread pattern of structural co-modulation. The network of HC>SZ appears particularly dense, indicating a high degree of structural integration. We observed strong co-modulation between the cognitive control and visual networks. In contrast, the network of SZ>HC showed significant co-modulation between the default mode and cognitive control networks.

### 3.2. Co-modulation differences for cognitive scores (controls and schizophrenia)

To understand the prospective clinical relevance of structural co-modulation, we correlated the subject-specific matrices with the composite CMINDS cognitive score. This analysis was performed separately for HC and SZ, and a direct comparison was computed to assess group differences in brain-cognition relationships (Figure 4a). In the healthy control group, higher cognitive performance was linked mainly to increased co-modulation among the somatomotor, visual, and subcortical networks. Higher cognitive performance was also associated with decreased co-modulation between the cerebellum and cognitive control networks. The auditory network co-modulation patterns showed no significant relationship with cognitive scores when comparing healthy controls. Biologically, this suggests that in a healthy brain, strong structural connections within sensory and subcortical regions are essential for supporting high cognitive performance.

**Figure 4.**
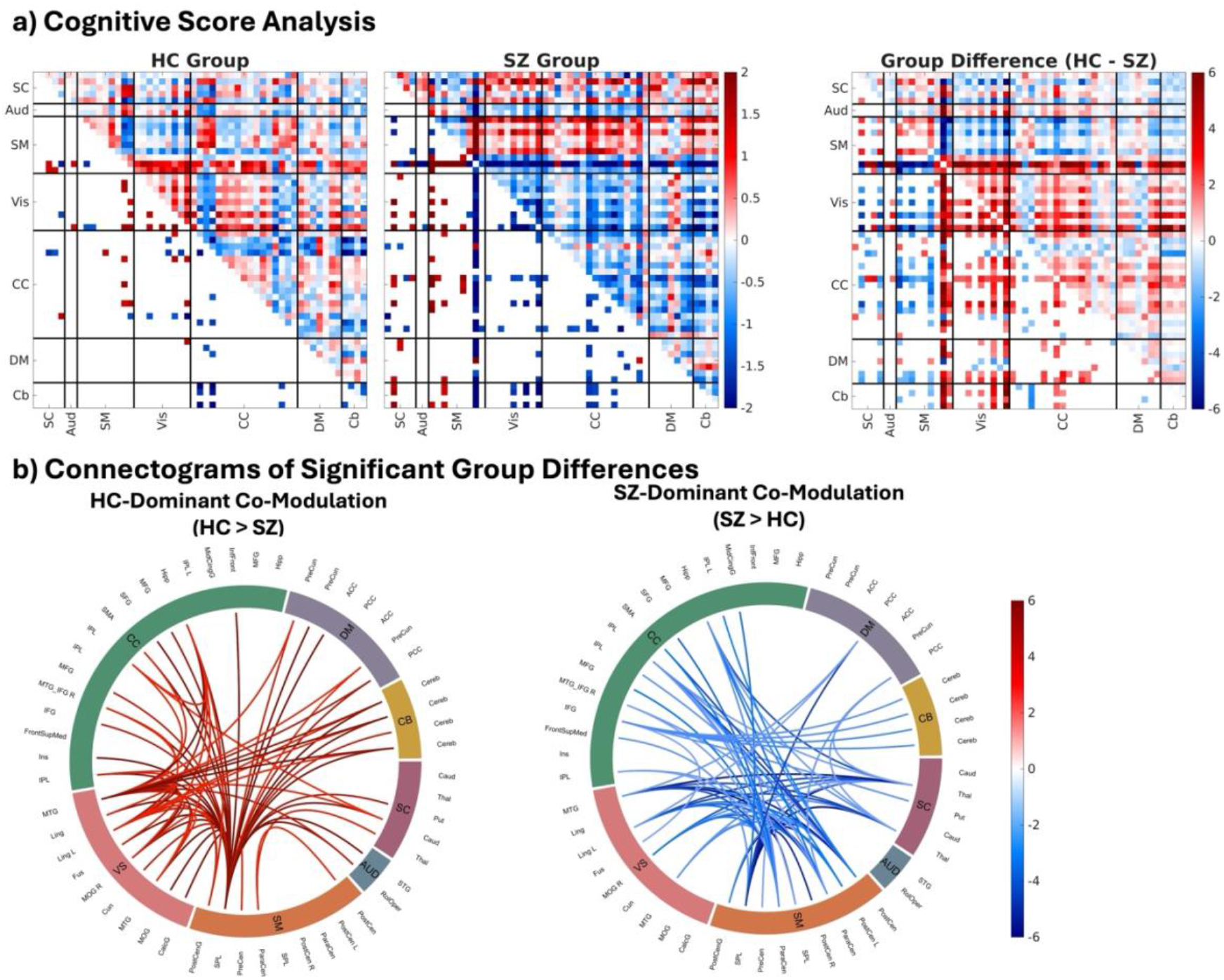
Co-modulation differences for cognitive scores. a. Co-modulation matrices display the signed log_10_(q) values from a whole-brain Pearson correlation between co-modulation values and a composite cognitive score. The two matrices on the left shows the correlation for the HC group and SZ group, respectively. The right matrix shows the difference between the two correlation matrices. For both matrices, red values indicate a positive correlation (higher co-modulation, better cognitive performance) and blue values indicate a negative correlation (lower co-modulation, better cognitive performance). The lower triangle displays only correlations that survived FDR correction (q < 0.05). b. Connectogram illustrating the top 5% most significant component associations. To visualize both effect direction and statistical significance, values are shown as signed log_10_q (FDR corrected) values. On this scale, we separated by the direction of the contrast. The left panel displays connections where the brain-cognition relationship was significantly more positive in controls (HC > SZ), while the right panel displays connections where the relationship was significantly more positive (or less negative) in patients (SZ > HC).

The schizophrenia patient group showed a markedly different pattern. The most prominent feature was a strong, statistically significant negative correlation emerging from a specific somatomotor component and its connections to nearly all other brain domains (including Subcortical, Auditory, and Visual). This indicates that in the SZ group, unlike controls, a different brain-cognition relationship emerges, where higher co-modulation involving this motor component is strongly related to poorer cognitive performance.

To directly test this apparent disruption, we analyzed the difference between the two correlation matrices (HC-SZ). This analysis revealed widespread, statistically significant group differences. In the visual networks, we observed a significantly more positive association in HC compared to SZ, indicating that coordinated structural patterns in these regions are beneficial for cognitive performance in healthy individuals, but this relationship is diminished or reversed in patients. Conversely, differences driven by the SZ group (SZ > HC) were found mainly in cognitive control and somatomotor network connections, reflecting the unique negative associations observed in schizophrenia. This suggests altered or compensatory structure– cognition relationships in schizophrenia.

To better visualize the spatial distribution and network topology of these widespread group differences, we rendered the significant findings from the HC-SZ matrix as a circular connectogram (Figure 4b). These plots are separated by the direction of the difference for clarity; the left panel shows connections where the correlation was significantly more positive in HC (HC > SZ), and the right panel shows connections where the correlation was significantly more positive (or less negative) in SZ (SZ > HC). All connections shown are statistically significant (q < 0.05), with the line color by the signed log_10_(q) values to represent the magnitude of the group difference. We found that positive brain-cognition relationships in controls are particularly concentrated within the visual and cognitive control networks. In patients, it is possible we found the the somatomotor networks were a critical locus of altered co-modulation relationships.

### 3.3. Co-modulation links to symptoms

The symptom analysis revealed distinct patterns for positive and negative symptoms (Figure 5). For negative symptoms, the matrix displays the signed log_10_(p) values; we observed visually apparent trends in the uncorrected data (upper triangle). The most distinct pattern was linked to the motor network, which showed widespread negative correlations within its own. This trend suggests that lower co-modulation involving the motor network is associated with higher negative symptom severity.

**Figure 5.**
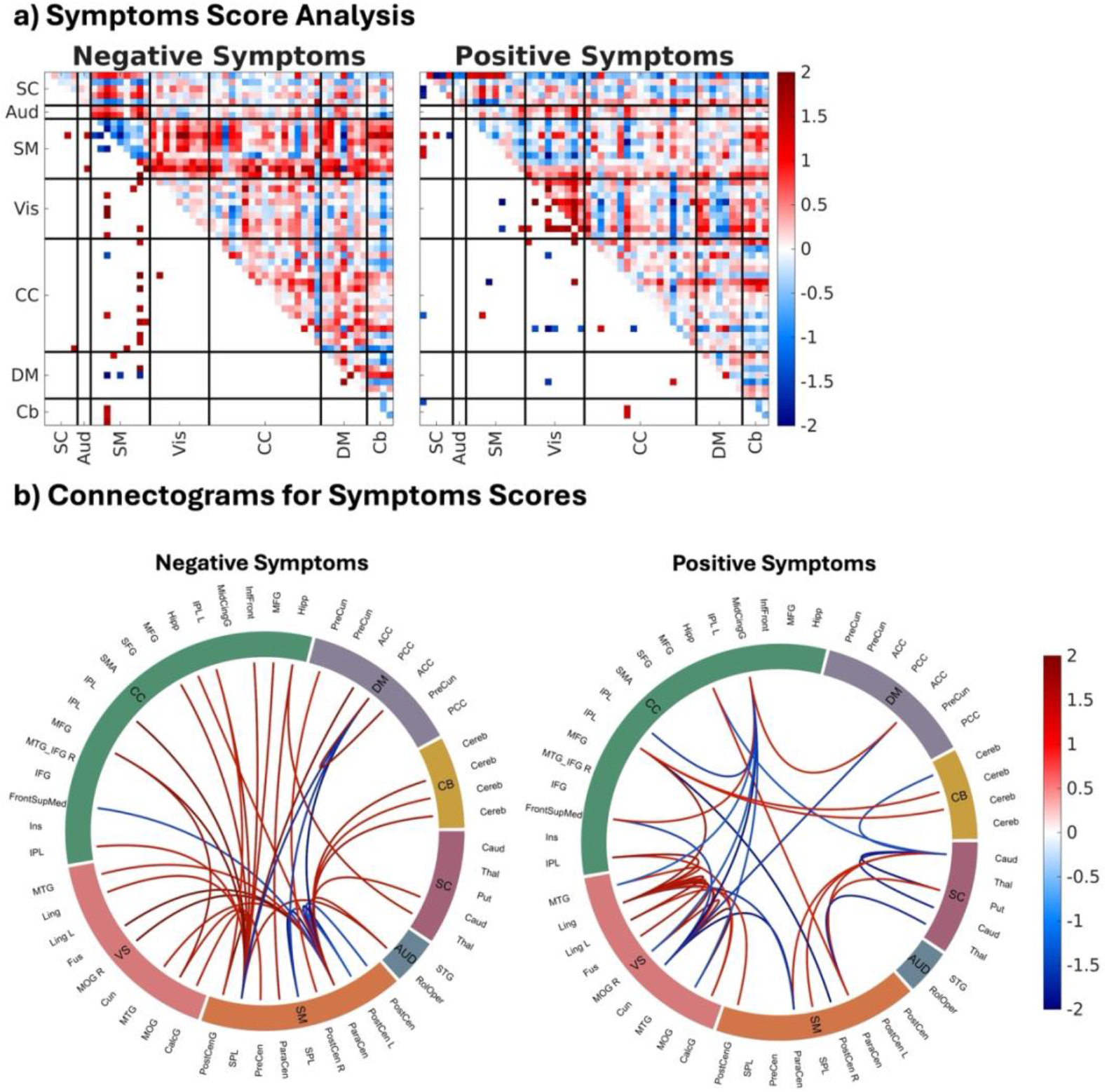
Co-modulation analysis for symptoms differences. a. Co-modulation matrices display the signed log_10_(p) values from a whole-brain Pearson correlation between co-modulation values and PANSS scores. The left matrix shows the correlation with negative symptoms, and the right matrix shows the correlation with positive symptoms. For both matrices, red values indicate a positive correlation (higher co-modulation, higher symptom severity) and blue values indicate a negative correlation (lower co-modulation, higher symptom severity). The lower triangle displays only correlations that survived FDR correction (p < 0.05). b. Connectogram illustrating significant component associations with symptoms scores (all connections p < 0.05). Results are plotted separately for negative symptoms (left) and positive symptoms (right). To visualize both effect direction and statistical significance, values are shown as signed log_10_p (FDR corrected) values. Positive values (red) indicate a positive correlation (higher co-modulation linked to higher symptoms), and negative values (blue) indicate a negative correlation (higher co-modulation linked to lower symptoms).

As shown in the lower triangle, a few of these key associations remained statistically significant even after a stringent FDR correction for multiple comparisons. This is a particularly noteworthy finding, given that clinical symptom profiles are notoriously difficult to map. The fact that this method can detect clear, structured, and statistically robust associations suggests that co-modulation holds promise for future clinical application. These findings should be interpreted with this in mind, and with the caution that results may be limited by unmeasured variables, such as the potential confounding effects of antipsychotic medication on brain structure.

For positive symptoms, one of the notable trends was a pattern of positive correlations within the visual domain. While these trends did not survive FDR correction, they suggest that larger studies might reveal positive correlations between intra-visual structural co-modulation and positive symptom severity, a finding that aligns with the prevalence of visual processing abnormalities in schizophrenia. This finding underscores the well-known challenge in mapping heterogeneous symptoms like hallucinations and delusions to specific structural patterns (Butler and Javitt, 2005, Butler et al., 2008).

The connectograms in Figure 5b are presented to provide a clear illustration of these associations. The plot for negative symptoms (left panel) revelas the dominant pattern from the matrix analysis, highlighting the somatomotor networks as a key source of widespread negative and positive correlations. The plot for positive symptoms (right panel) shows a distinct cluster of elements primarily between structural features belonging to the visual domain.

## 4. Discussion

This study introduces co-modulation, a novel method to quantify the inter-relationship between structural networks derived from subject-specific SBM data. This approach represents a critical advance over traditional SBM, which typically focuses on the magnitude of gray matter changes within a given component (Kotoski et al., 2024a), overlooking the variation among them. By capturing these inter-network dependencies, the co-modulation framework provides a new approach to investigate whether neuropathology in disorders like schizophrenia extends beyond localized structural deficits to a broader breakdown in structural organization. This method provides a way not only to map which structural features are affected but also understand how distributed brain systems are systematically related at the individual level.

The application of this method revealed significant alterations of structural co-modulation pattern in schizophrenia. Our primary finding was a widespread reduction in structural co-modulation in patients compared to healthy controls (Figure 3). This supported the hypothesis that schizophrenia is a disorder of failed network integration (van Erp et al., 2018). Rather than being randomly organized, these reductions were not random, they were most prominent within and between higher-order networks, including the visual, and cognitive control. Specifically, we found a profound decrease of co-modulation in patients centered on the right middle occipital gyrus. These key dorsal visual stream regions, which were strongly co-modulated with executive control and attentional regions in the frontal lobe of healthy controls, exhibited significantly decreased covariance in patients. This finding provides a specific structural basis for the well-known breakdown between visual-spatial attention and executive function in schizophrenia (Butler et al., 2008).

However, this pattern was not a simple, uniform loss. We also observed specific increases in co-modulation in patients involving the default mode network and parts of the cerebellum. This novel insight suggests that schizophrenia may be characterized not just by a de-coordination of structural architecture, but by a significant reorganization of it (Guo et al., 2015, Wang et al., 2014). The demeaned matrices visually confirmed this (Figure 3c), showing that while controls had stronger than average co-modulation in key cognitive domains, patients exhibited a mixed pattern of both weaker than average co-modulation and these specific focal increases. This reorganization was most evident in two specific patterns. First, we observed an increase in co-modulation centered on the hippocampus, which reflected structural coupling with primary somatosensory and visual cortices. This finding aligns with the hypothesis of a breakdown in the brain’s ability to filter relevant from irrelevant stimuli, potentially causing raw sensory input to be incorrectly integrated into memory systems (Ford et al., 2015, Kapur, 2003, Lodge and Grace, 2011). This structural configuration provides a potential anatomical substrate for aberrant salience, which is the state where neutral events or objects feel intensely and personally significant (De Pieri et al., 2025). Second, we found a strong connection between the anterior cingulate cortex (ACC) and the cerebellum. The ACC plays a key role in the brain’s error-detection system (Carter et al., 1998), and the cerebellum helps coordinate and fine-tune complex thoughts (Botvinick et al., 2001, Hirsch et al., 2025). This pattern of excessive structural covariance may reflect a static network architecture that creates a vulnerability for the cognitive rigidity or disorganized thinking often observed in schizophrenia (Andreasen et al., 1999).

We also demonstrated that these co-modulation patterns are functionally and clinically relevant. We found that the relationship between structural organization and cognitive performance was different between groups (Figure 4). In healthy controls, we found a characteristic pattern of healthy structural organization: cognitive performance was strongly linked to the co-modulation of a single component, the superior parietal lobe. This key dorsal attention network hub (Corbetta and Shulman, 2002) was integrated with visual, frontal, and default-mode networks, suggesting that cognitive performance in healthy individuals relies on the integration of spatial attention with other high-order systems.

This relationship was disrupted in schizophrenia. Patients exhibited distinct and prospectively maladaptive brain-cognition relationships. The attentional-cognitive link was absent, and cognitive performance in patients was instead strongly linked to the co-modulation of primary sensorimotor regions and the thalamus, suggesting a shift to a pattern of structural organization between lower-level or primary order regions that was related to poorer cognitive performance. This finding of an altered structure-cognition relationship in schizophrenia suggests that the organizational breakdown has direct functional consequences. This concept is strongly supported by recent work using person-based similarity index (PBSI) research which similarly shows that individuals with schizophrenia exhibit greater deviation from a healthy structural brain norm, and that magnitudes of this individual-level deviation are linked to cognitive and clinical severity (Jo et al., 2025).

While our investigation of brain-symptom relationships is preliminary, they nevertheless revealed clinically interpretable patterns (Figure 5). For example, we observed an association between higher negative symptom severity and lower co-modulation within the somatomotor network. This finding aligns with growing evidence that abnormalities in the brain’s motor system contribute to core negative symptoms, such as avolition and psychomotor slowing (Mehmood et al., 2026, Walther and Strik, 2012). We also found that higher intra-visual co-modulation was linked to greater positive symptom severity, an association that is highly consistent with the prevalence of visual processing abnormalities and hallucinations in schizophrenia (Butler and Javitt, 2005, Tan et al., 2025). However, the fact that these anatomically specific associations did not withstand correction for multiple comparisons is not unexpected. It shows the profound challenge of mapping heterogeneous clinical profiles onto specific neural substrates, especially given potential confounding variables such as antipsychotic medication, which were not accounted for in this analysis. The ability of our co-modulation method to detect these structured patterns at all, is promising. It suggests this method holds a degree of sensitivity to subtle clinical-anatomical relationships that warrants future investigation, perhaps in larger, medication-naive, or first-episode cohorts where statistical power may be greater and confounds are minimized.

In sum, this study introduces structural co-modulation as a novel and valuable method for quantifying the organization of brain structure. We demonstrated its utility in capturing structural brain alterations in schizophrenia, revealing a global reduction in the coordination between structural networks, further accumulating evidence for the hypothesis that distributed brain alterations are a key underlying feature of the disorder. We also demonstrated that co-modulation patterns have behavioral relevance, finding that the healthy relationship between structural organization and cognition is fundamentally disrupted in schizophrenia. In conclusion, our work provides new evidence that schizophrenia is a disorder of large-scale structural discoordination and establishes the co-modulation framework as a promising tool for understanding complex gray matter relationships and for mapping brain pathologies.

